# MarkerScout: A Disease-Agnostic Machine Learning Framework for Biomarker Prediction from Multi-Scale Mechanistic Models

**DOI:** 10.64898/2026.05.30.727266

**Authors:** Robert Moore, Frank Agayie-Ntim, Lindsey B. Crawford, Prakash Packrisamy, Ahmed Abdeen Hamed, Tomáš Helikar

## Abstract

Identifying robust biomarkers from high-dimensional biomedical data is a central challenge in translational research, but candidate rankings produced by any single feature-selection or classification method depend on algorithmic choices and rarely reproduce across pipelines. We present a disease-agnostic machine-learning framework that addresses this dependence by systematically benchmarking 25 (feature-selection x classifier) pipelines under five-fold stratified cross-validation, aggregating per-feature evidence by two independent methods (a weighted-selection consensus score and Robust Rank Aggregation), and characterizing the direction of each candidate using Cohen’s *d*. We demonstrate the framework on immune-response measurements from two clinical phases: SARS-CoV-2 hospitalization and intensive-care admission; obtaining cross-validated mean F1 above 0.99 with balanced classification errors and producing tiered, direction-aware biomarker lists per phase. Interleukin-18 (IL-18) reached the strongest tier in both phases with consistent direction. The framework generalizes to any binary clinical classification problem and supports principled, reproducible biomarker prioritization.

## INTRODUCTION

Biomarkers are now a cornerstone of modern translational research, enabling earlier diagnosis, deeper mechanistic insights, and precise treatments for highly varied diseases. They act as measurable indicators of biological states by capturing the molecular, cellular, and physiological processes that drive disease progression and how a patient responds to therapy. Recent studies show how different biological mechanisms, from immune imbalances to metabolic changes, can be identified through molecular signatures that improve clinical decisions. For example, in acute myocardial infarction (AMI), Miao et al. identified 6 genes associated with cuproptosis and ferroptosis, with expression patterns that distinguished patients from healthy controls. These markers diagnosed AMI with high sensitivity and specificity, showing how multi-omic biomarkers can reveal previously hidden disease mechanisms ^1^. Similarly, Grywalska et al. found that higher immune-checkpoint expression and pro-inflammatory cytokines are effective biomarkers in pediatric vulvar lichen sclerosus, illustrating how immune pathways can be used for early detection in chronic inflammatory disorders ^2^. Together, these findings demonstrate that biomarkers do more than just classify a disease; they reveal the immunological, metabolic, and regulatory circuits that drive pathology, making them essential tools for precision medicine.

### 2. Machine Learning: A Driver of Biological Discovery

Machine learning has become a transformative force in biological discovery. It allows researchers to uncover meaningful structures, mechanistic insights, and predictive patterns within large, complex datasets. Early foundational work showed that supervised and unsupervised learning could effectively classify biological states, infer regulatory relationships, and integrate various omics modalities, building the methodological core of today’s computational biology (Tarca et al., 2007; Lee, 2009; Ghosh & Dasgupta, 2022)^3–5^. As data grew more complex, new frameworks emerged to format, curate, and model high-dimensional molecular information, ensuring that machine-learning algorithms could work effectively with genomic, proteomic, structural, and network-level data (Duran-Frigola et al., 2019; Auslander et al., 2021; Liu & Zhang, 2024)^6–8^. Deep learning further pushed this boundary by enabling representation learning from raw sequences, images, and single-cell profiles, while advances in large-scale ML frameworks provided the necessary computational power (Ching et al., 2018; Nguyen et al., 2019; Goshisht, 2024)^9–11^. Beyond simple prediction, machine learning now drives discovery through network-based and knowledge-guided approaches, which help identify drug indications, reconstruct regulatory circuits, and integrate mechanistic knowledge into predictive models (Gilvary et al., 2020; Erbe et al., 2022; Karpatne et al., 2022)^12–14^. Deep generative models take this even further by learning latent biological structures and proposing novel molecular configurations (Lopez et al., 2020)^15^. Ultimately, machine learning is more than just a tool; it is an essential lens that reveals the hidden architecture of biological systems and makes complex experimental questions much more approachable.

### 3. Applying Machine Learning to Biomarker Discovery

Machine learning can detect the subtle, multivariate, and context-dependent patterns needed to identify reliable signatures of disease and therapy response. While traditional statistical methods often struggle with the noise and high dimensionality of omics data, machine-learning methods can extract stable, interpretable features. For instance, Hédou et al. developed Stabl, a framework that finds reproducible biomarkers by ensuring stability across resampling, showing that clear signals can be recovered even from noisy clinical data^16^. Explainable models have also helped bridge the gap between prediction and actual biological mechanisms. Ijaz et al., for example, used interpretable models to find immune-inflammatory biomarkers for drug-resistant epilepsy^17^. Integrating multiple omics data types is another powerful approach; Li et al. showed that tri-omics frameworks can identify biomarkers for sepsis, revealing how immune and metabolic systems fail together during critical illness^18^. Other studies have applied ML to immune signatures in joint infections, polycystic ovarian syndrome, and immune-checkpoint pathways like B7x, which moved from discovery toward clinical trials^19^. Deep neural networks are also helping predict how patients will respond to immunotherapies. These collective advances show that machine-learning-driven discovery offers a unified way to understand disease, group patients effectively, and personalize care across many medical fields.

We present a flexible, disease-agnostic machine-learning pipeline that works across diverse patient groups and clinical stages. The workflow starts by bringing together different datasets and outcome labels into a single preprocessing and feature-selection layer, ensuring all models use harmonized data. Building on this, the pipeline tests various model types and pairing strategies in parallel, using a grid-search engine to find the most reliable options for each disease stage. Once the best models are chosen, we use specific aggregation strategies, such as consolidation for hospitalization data and consensus-frequency scoring for ICU cases, to create stable biomarker signatures. This setup allows one reusable framework to handle changing disease dynamics while keeping specific biological signals intact, ultimately making biomarker discovery more reproducible and robust.

## RESULTS

### Top-Ranked Classification Pipelines for SARS-CoV-2 Hospitalization and ICU Cohorts

For each of the two SARS-CoV-2 clinical-phase cohorts: hospitalization (COV-HOSP) and intensive-care (COV-ICU), we benchmarked the full factorial of five feature-selection algorithms (L1Selection, MutualInformation, RFE_LinearSVM, SelectKBest, VIFFiltering) and five binary classifiers (LogisticRegression, MLPClassifier, kNN, DecisionTreeClassifier, GaussianNB), yielding 25 distinct (Selector × Classifier) pipelines per cohort and 50 pipelines in aggregate. Each pipeline was evaluated using a 5-fold stratified cross-validation scheme against the binary “Clearance” outcome (cleared = 0, not_cleared = 1). Metrics were computed independently on each fold and then averaged to obtain the mean F1, recall, precision, and accuracy values reported below (Table 1).

**Table 1.** Top 10 (Selector × Classifier) pipelines per cohort, ranked by mean F1.

| Cohort | Rank | Feature-Selection<br>Method | Classifier | Mean F1<br>(SD) | Mean<br>Recall | Mean<br>Precision | Mean<br>Accuracy |
| --- | --- | --- | --- | --- | --- | --- | --- |
| COV-HOSP | 1 | RFE_LinearSVM | LogisticRegression | 0.9916<br>(0.0013) | 0.9918 | 0.9914 | 0.9982 |
| COV-HOSP | 2 | L1Selection | LogisticRegression | 0.9905<br>(0.0022) | 0.9909 | 0.9901 | 0.9979 |
| COV-HOSP | 3 | RFE_LinearSVM | MLPClassifier | 0.9895<br>(0.0024) | 0.9912 | 0.9881 | 0.9977 |
| COV-HOSP | 4 | L1Selection | MLPClassifier | 0.9893<br>(0.0013) | 0.9895 | 0.9892 | 0.9977 |
| COV-HOSP | 5 | RFE_LinearSVM | kNN | 0.9882<br>(0.0016) | 0.9877 | 0.9887 | 0.9975 |
| COV-HOSP | 6 | RFE_LinearSVM | DecisionTreeClassifier | 0.9866<br>(0.0015) | 0.9870 | 0.9862 | 0.9971 |
| COV-HOSP | 7 | L1Selection | DecisionTreeClassifier | 0.9848<br>(0.0012) | 0.9846 | 0.9849 | 0.9967 |
| COV-HOSP | 8 | SelectKBest | LogisticRegression | 0.9848<br>(0.0020) | 0.9844 | 0.9852 | 0.9967 |
| COV-HOSP | 9 | SelectKBest | MLPClassifier | 0.9843<br>(0.0038) | 0.9829 | 0.9862 | 0.9967 |
| COV-HOSP | 10 | L1Selection | kNN | 0.9836<br>(0.0030) | 0.9845 | 0.9827 | 0.9965 |
| COV-ICU | 1 | L1Selection | LogisticRegression | 0.9941<br>(0.0021) | 0.9940 | 0.9941 | 0.9965 |
| COV-ICU | 2 | L1Selection | MLPClassifier | 0.9934<br>(0.0031) | 0.9931 | 0.9939 | 0.9961 |
| COV-ICU | 3 | MutualInformation | LogisticRegression | 0.9932<br>(0.0027) | 0.9930 | 0.9934 | 0.9959 |
| COV-ICU | 4 | SelectKBest | LogisticRegression | 0.9926<br>(0.0021) | 0.9924 | 0.9928 | 0.9956 |
| COV-ICU | 5 | RFE_LinearSVM | LogisticRegression | 0.9914<br>(0.0030) | 0.9908 | 0.9920 | 0.9949 |
| COV-ICU | 6 | RFE_LinearSVM | MLPClassifier | 0.9914<br>(0.0031) | 0.9912 | 0.9916 | 0.9949 |
| COV-ICU | 7 | MutualInformation | MLPClassifier | 0.9911<br>(0.0031) | 0.9900 | 0.9922 | 0.9947 |
| COV-ICU | 8 | SelectKBest | MLPClassifier | 0.9908<br>(0.0030) | 0.9898 | 0.9917 | 0.9945 |
| COV-ICU | 9 | SelectKBest | kNN | 0.9825<br>(0.0025) | 0.9815 | 0.9835 | 0.9896 |
| COV-ICU | 10 | MutualInformation | kNN | 0.9822<br>(0.0024) | 0.9814 | 0.9830 | 0.9894 |

All values are mean ± standard deviation across 5 stratified folds. “Cleared” = 0, “Not cleared” = 1 in the binary Clearance target.

In both cohorts the top-ranked pipelines cluster within a narrow, high-performing band of mean F1 scores. In COV-HOSP, the top 10 mean F1 scores fall in the range 0.9836 – 0.9916, while in COV-ICU they fall in 0.9822 – 0.9941, which is a top-10 spread of only 0.008 F1 units in COV-HOSP and 0.012 in COV-ICU. The single best pipeline in COV-HOSP is RFE_LinearSVM paired with LogisticRegression (mean F1 = 0.9916 ± 0.0013, mean accuracy = 0.9982), and the single best pipeline in COV-ICU is L1Selection paired with LogisticRegression (mean F1 = 0.9941 ± 0.0021, mean accuracy = 0.9965). Across all 20 top-10 rows, the cross-fold standard deviation of F1 never exceeds 0.004, and precision and recall are tightly balanced (absolute gap ≤ 0.005 within every row), indicating independence of each experiment and that the pipeline avoids the algorithmic selection bias by successfully identifying the best runs. Figure 2 presents a heatmap of the top-10 performing selector-classifier algorithms.

**Figure 1.**
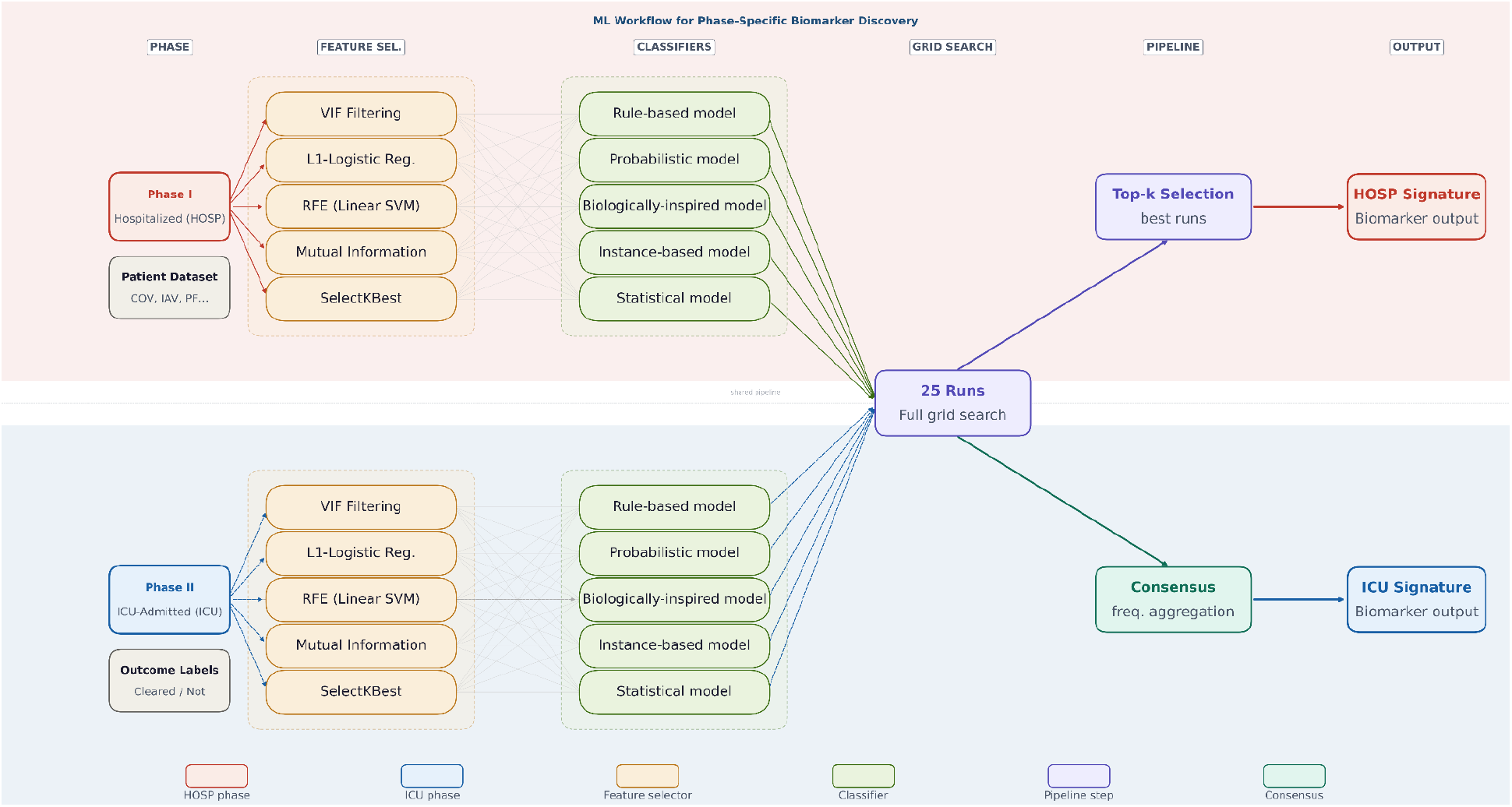
Adaptive, disease-agnostic machine-learning workflow for phase-specific biomarker discovery. This pipeline starts with a preprocessing and feature-selection layer to harmonize diverse datasets and outcome labels. It systematically benchmarks 25 (feature-selection × classifier) pipelines using stratified 5-fold cross-validation. Stable biomarker signatures are created by aggregating per-feature evidence using two independent methods: a weighted-selection consensus score and Robust Rank Aggregation. Finally, the framework characterizes the direction of each candidate using Cohen’s d.

**Figure 2.**
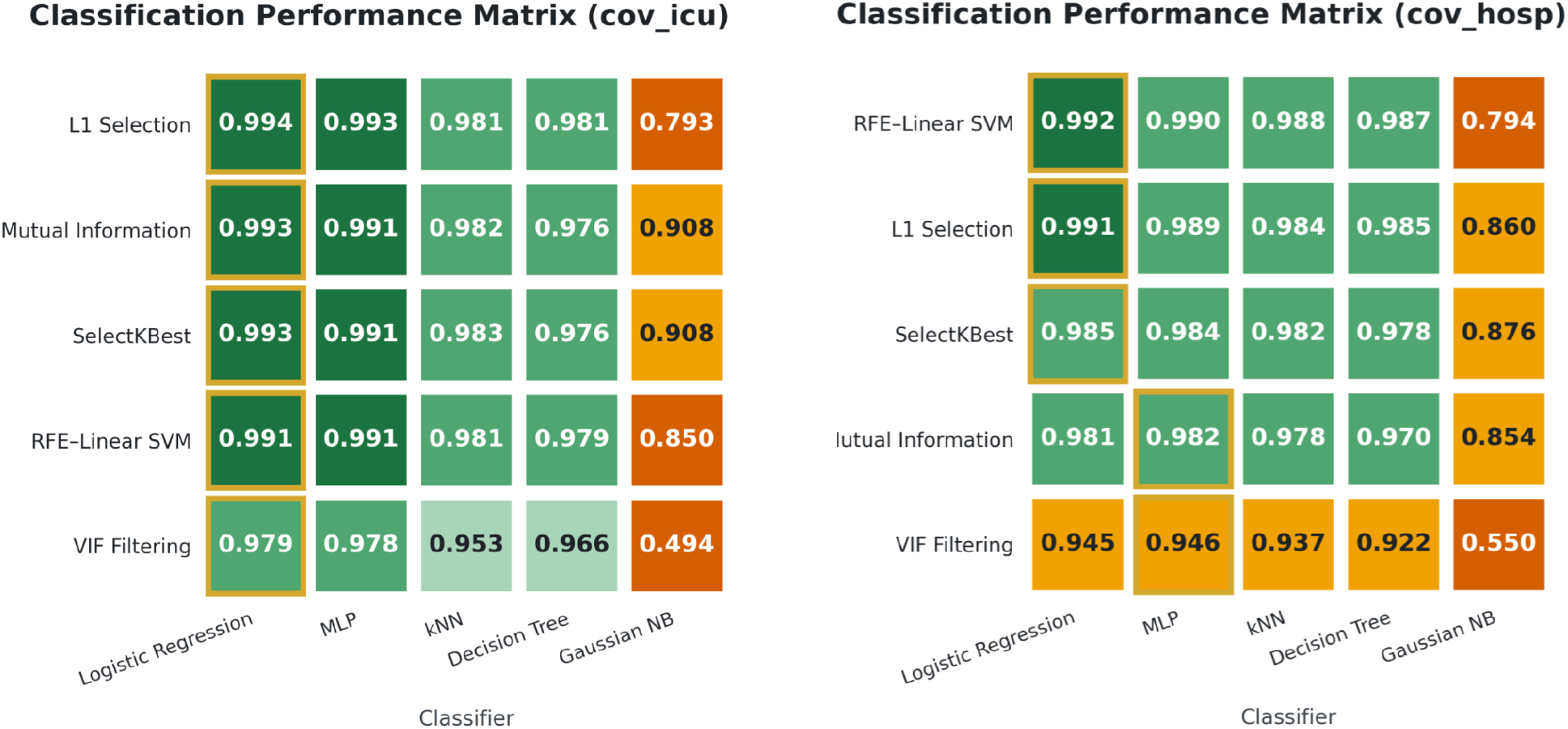
Top-10 selector–classifier pipelines for COV-HOSP and COV-ICU. Both cohorts show tightly clustered high-performance bands: 0.9836–0.9916 (COV-HOSP) and 0.9822–0.9941 (COV-ICU). The best pipelines are RFE_LinearSVM + LogisticRegression for COV-HOSP and L1Selection + LogisticRegression for COV-ICU. Cross-fold F1 variability remains < 0.004, with precision–recall gaps ≤ 0.005, indicating stable, unbiased performance. The heatmap displays the top-10 pipelines for each cohort.

### Error profile of the top-10 pipelines — COV-HOSP and COV-ICU

Per-fold false-positive (FP) and false-negative (FN) counts for the ten highest-ranked (Selector × Classifier) pipelines in each cohort, under 5-fold stratified cross-validation against the binary Clearance outcome (cleared = 0, not-cleared = 1). Means and standard deviations are computed across the 5 folds; total FP and total FN are the summed counts across all folds.

The observations in Tables 2a and 2b show that, in both cohorts, the rank-1 pipeline simultaneously minimises mean false-positive (FP) and mean false-negative (FN) counts across the top-10 — RFE_LinearSVM + LogisticRegression in COV-HOSP (mean FP = 18.6 ± 3.3, mean FN = 17.4 ± 4.9) and L1Selection + LogisticRegression in COV-ICU (mean FP = 2.0 ± 1.1, mean FN = 2.0 ± 1.3) — so the F1 leader is also the joint error leader, with no top-10 configuration preferable on either error axis. Across the top-10 of both cohorts, FP and FN counts remained balanced (total FP / FN ratio 0.81 – 1.46 in COV-HOSP, 0.81 – 1.46 in COV-ICU), indicating that the classifiers do not preferentially over- or under-predict the not-cleared class. Absolute error counts scaled with cohort size (top-1 totals 180 errors in COV-HOSP vs. 20 in COV-ICU across 5 folds, a ∼9× ratio consistent with the smaller COV-ICU test population), but the relative ordering and the rank-1 dominance pattern were preserved across cohorts.

**Table 2a.**
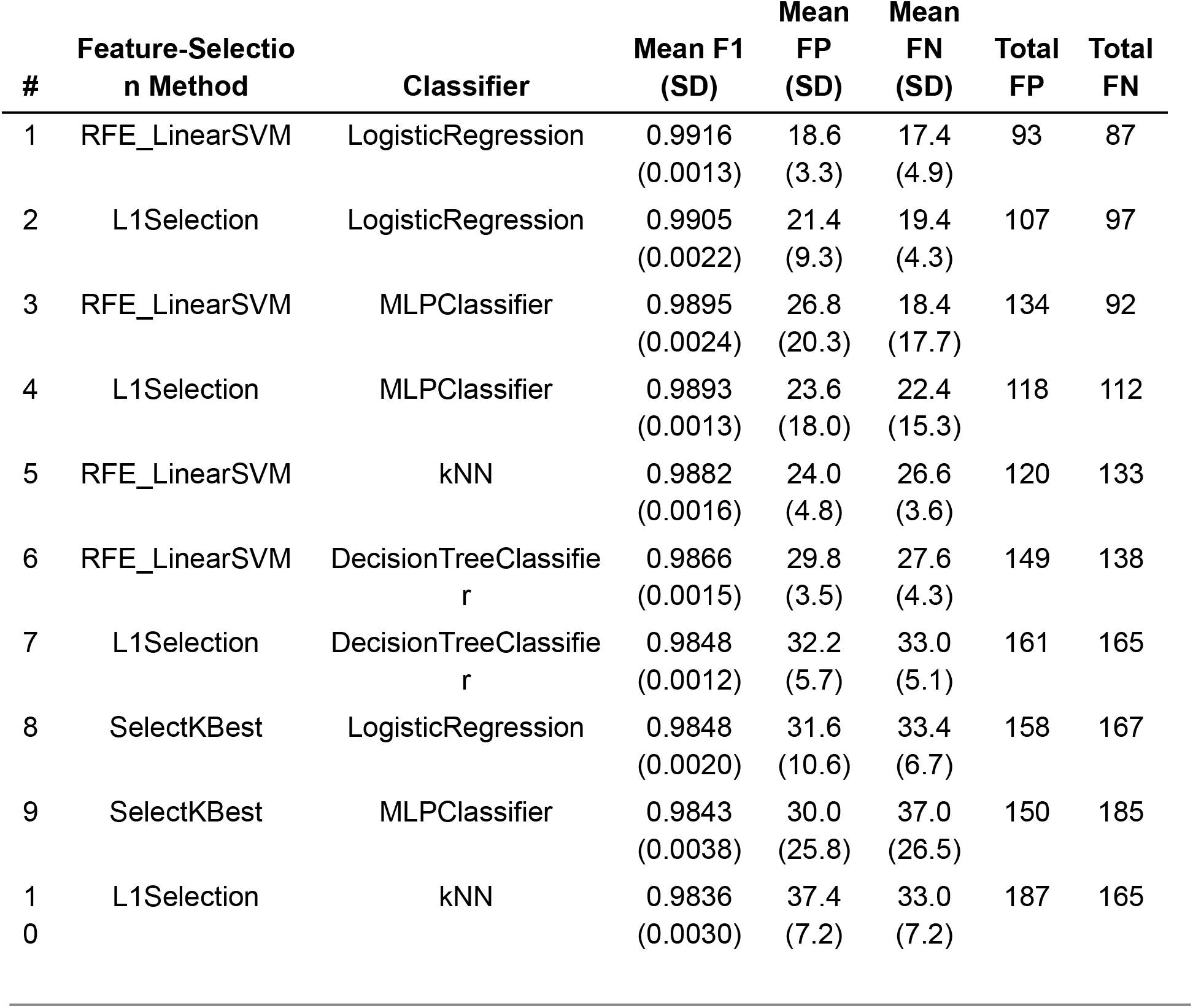
COV-HOSP: error profile of the top-10 pipelines.

| # | Feature-Selection Method | Classifier | Mean F1 (SD) | Mean FP (SD) | Mean FN (SD) | Total FP | Total FN |
| --- | --- | --- | --- | --- | --- | --- | --- |
| 1 | RFE_LinearSVM | LogisticRegression | 0.9916<br>(0.0013) | 18.6<br>(3.3) | 17.4<br>(4.9) | 93 | 87 |
| 2 | L1Selection | LogisticRegression | 0.9905<br>(0.0022) | 21.4<br>(9.3) | 19.4<br>(4.3) | 107 | 97 |
| 3 | RFE_LinearSVM | MLPClassifier | 0.9895<br>(0.0024) | 26.8<br>(20.3) | 18.4<br>(17.7) | 134 | 92 |
| 4 | L1Selection | MLPClassifier | 0.9893<br>(0.0013) | 23.6<br>(18.0) | 22.4<br>(15.3) | 118 | 112 |
| 5 | RFE_LinearSVM | kNN | 0.9882<br>(0.0016) | 24.0<br>(4.8) | 26.6<br>(3.6) | 120 | 133 |
| 6 | RFE_LinearSVM | DecisionTreeClassifier | 0.9866<br>(0.0015) | 29.8<br>(3.5) | 27.6<br>(4.3) | 149 | 138 |
| 7 | L1Selection | DecisionTreeClassifier | 0.9848<br>(0.0012) | 32.2<br>(5.7) | 33.0<br>(5.1) | 161 | 165 |
| 8 | SelectKBest | LogisticRegression | 0.9848<br>(0.0020) | 31.6<br>(10.6) | 33.4<br>(6.7) | 158 | 167 |
| 9 | SelectKBest | MLPClassifier | 0.9843<br>(0.0038) | 30.0<br>(25.8) | 37.0<br>(26.5) | 150 | 185 |
| 10 | L1Selection | kNN | 0.9836<br>(0.0030) | 37.4<br>(7.2) | 33.0<br>(7.2) | 187 | 165 |

**Table 2b.** COV-ICU: error profile of the top-10 pipelines.

| # | Feature-Selection<br>Method | Classifier | Mean F1<br>(SD) | Mean<br>FP<br>(SD) | Mean<br>FN<br>(SD) | Total<br>FP | Total<br>FN |
| --- | --- | --- | --- | --- | --- | --- | --- |
| 1 | L1Selection | LogisticRegression | 0.9941<br>(0.0021) | 2.0<br>(1.1) | 2.0<br>(1.3) | 10 | 10 |
| 2 | L1Selection | MLPClassifier | 0.9934<br>(0.0031) | 2.0<br>(0.6) | 2.4<br>(2.2) | 10 | 12 |
| 3 | MutualInformation | LogisticRegression | 0.9932<br>(0.0027) | 2.2<br>(1.5) | 2.4<br>(1.9) | 11 | 12 |
| 4 | SelectKBest | LogisticRegression | 0.9926<br>(0.0021) | 2.4<br>(1.5) | 2.6<br>(1.6) | 12 | 13 |
| 5 | RFE_LinearSVM | LogisticRegression | 0.9914<br>(0.0030) | 2.6<br>(1.5) | 3.2<br>(2.0) | 13 | 16 |
| 6 | RFE_LinearSVM | MLPClassifier | 0.9914<br>(0.0031) | 2.8<br>(1.7) | 3.0<br>(2.1) | 14 | 15 |
| 7 | MutualInformation | MLPClassifier | 0.9911<br>(0.0031) | 2.4<br>(1.2) | 3.6<br>(1.9) | 12 | 18 |
| 8 | SelectKBest | MLPClassifier | 0.9908<br>(0.0030) | 2.6<br>(1.0) | 3.6<br>(1.9) | 13 | 18 |
| 9 | SelectKBest | kNN | 0.9825<br>(0.0025) | 5.4<br>(2.6) | 6.4<br>(2.1) | 27 | 32 |
| 10 | MutualInformation | kNN | 0.9822<br>(0.0024) | 5.6<br>(2.7) | 6.4<br>(2.1) | 28 | 32 |

### Feature scoring and tier assignment

To move from “which pipelines perform best” to “which features carry the signal,” we ranked features by two independent methods: Consensus (per-feature Σ of selection-frequency × mean F1 across all 25 pipelines) and Robust Rank Aggregation (RRA) with a Bonferroni-corrected ρ. We then took the union of each method’s top 20. Features were placed into three tiers:

- **Tier 1 (Strong):** present in both top-20 lists with rank shift ≤ 2 and -log_10_(ρ_Bonferroni) ≥ 2.
- **Tier 2 (Moderate):** present in both top-20 lists but failing one Tier-1 criterion.
- **Tier 3 (Exploratory):** present in only one method’s top-20.

In COV-HOSP, the procedure yielded 3 Tier-1, 12 Tier-2, and 12 Tier-3 features (25 total), with IL18_s, DC_mature, and IL10_s populating Tier 1. In COV-ICU, it yielded 8 Tier-1, 6 Tier-2, and 10 Tier-3 features (25 total), with GMCSF_s, IFNb_s, IL17_s, IL18_s, IL1B_s, and IFNa_s populating Tier 1. Tier-3 features reflected method-specific signals from both ranking approaches. In COV-HOSP, five Tier-3 features were RRA-only (DC_APC, DC_pDC, IL33_s, IFNg_s, ILC2), while five were Consensus-only (TGFb_s, TNFa_s, MCSF_s, Neutrophil_activated, Tcell_CD8_Naive). In COV-ICU, six Tier-3 features were RRA-only (DC_APC, IL1A_s, ILC2, IgM_s, NK_bright, NK_Dim), while six were Consensus-only (IL6_s, IgE_s, Neutrophil_activated, ROS, TGFb_s, TNFa_s). Across cohorts, IL18_s is the only feature reaching Tier 1 in both phases, making it the strongest cross-phase candidate discriminatory marker. IL17_s, IFNa_s, IFNb_s, and GMCSF_s are Tier-1 in COV-ICU and Tier-2 in COV-HOSP, supporting a robust but phase-modulated role. Figure 3 shows the outcome of the two ranking mechanisms for each of the cohorts.

**Figure 3.**
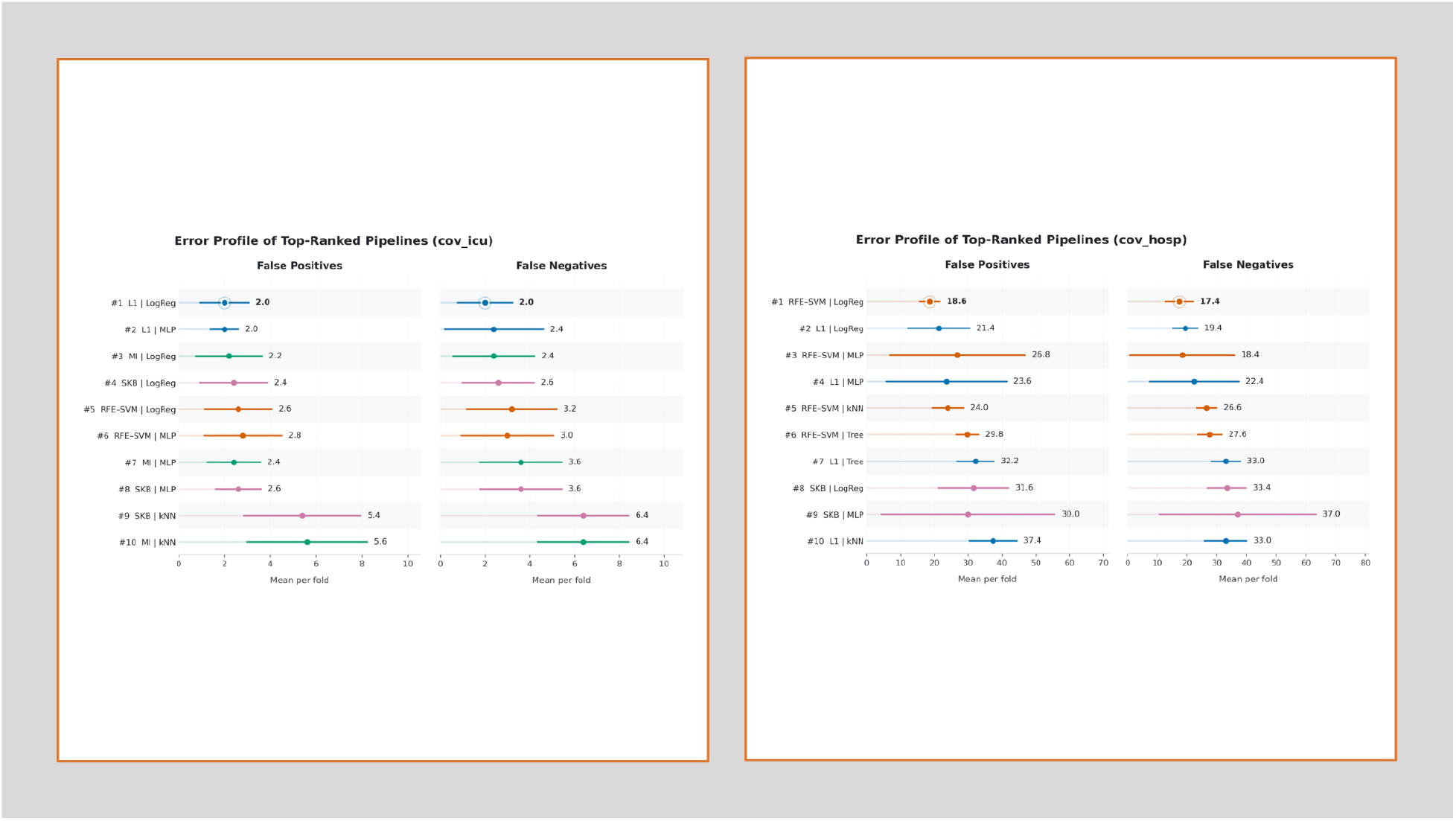
Error profiles of the top-10 selector–classifier pipelines in COV-HOSP and COV-ICU. Across both cohorts, the rank-1 pipeline minimizes mean false-positive (FP) and false-negative (FN) counts simultaneously. RFE_LinearSVM + LogisticRegression in COV-HOSP (mean FP = 18.6 ± 3.3; mean FN = 17.4 ± 4.9) and L1Selection + LogisticRegression in COV-ICU (mean FP = 2.0 ± 1.1; mean FN = 2.0 ± 1.3)—making the F1 leader also the joint error leader. FP and FN counts remain balanced across the top-10 (FP/FN ratio 0.81–1.46 in both cohorts), indicating no systematic bias toward over- or under-predicting the not-cleared class. Absolute error totals scale with cohort size (top-1 totals: 180 in COV-HOSP vs. 20 in COV-ICU across 5 folds), but the relative ordering and rank-1 dominance are preserved across cohorts.

### Direction of association using Cohen’s *d*

To characterize the direction of association independently of the ranking, we computed Cohen’s *d* for each tiered feature on the cleared versus not-cleared groups (a positive *d* indicates a higher mean in the not-cleared group, while a negative *d* indicates a higher mean in the cleared group). Among Tier-1 markers, the largest standardized effects in COV-HOSP were IL10_s (*d* = +2.04), IL18_s (*d* = −1.16), and DC_mature (*d* = −1.13); in COV-ICU, the largest effects were IL1B_s (*d* = +2.68), IFNa_s (*d* = +2.63), and IFNb_s (*d* = +2.57).

Ten features appeared on the tiered lists for both cohorts (COV-HOSP and COV-ICU), with their directions agreeing in 9 out of 10 cases. IL18_s, IL17_s, and GMCSF_s trended higher in cleared subjects in both cohorts (negative *d*), while IFNa_s, IFNb_s, IL10_s, IL12_s, IL9_s, and IFNg_s trended higher in not-cleared subjects (positive *d*). The only exception was DC_APC, which showed opposite Cohen’s *d* signs between cohorts (HOSP *d* = +0.04 versus ICU *d* = −0.58). Since the COV-HOSP magnitude is negligible (|*d*| < 0.1), this is considered a noise-level disagreement rather than a biological inversion. All five cross-phase candidate markers identified by the ranking analysis (IL18_s, IL17_s, IFNa_s, IFNb_s, GMCSF_s) consistently showed the same Cohen’s-*d* sign in both phases, reinforcing their cross-phase robustness. Table 2 presents the top features, their corresponding consensus and RRA ranking scores, and the direction. Figure 4 illustrates the Cohen’s *d* effect sizes for these tiered biomarkers, showing both the magnitude and direction of their associations in each cohort.

**Figure 4.**
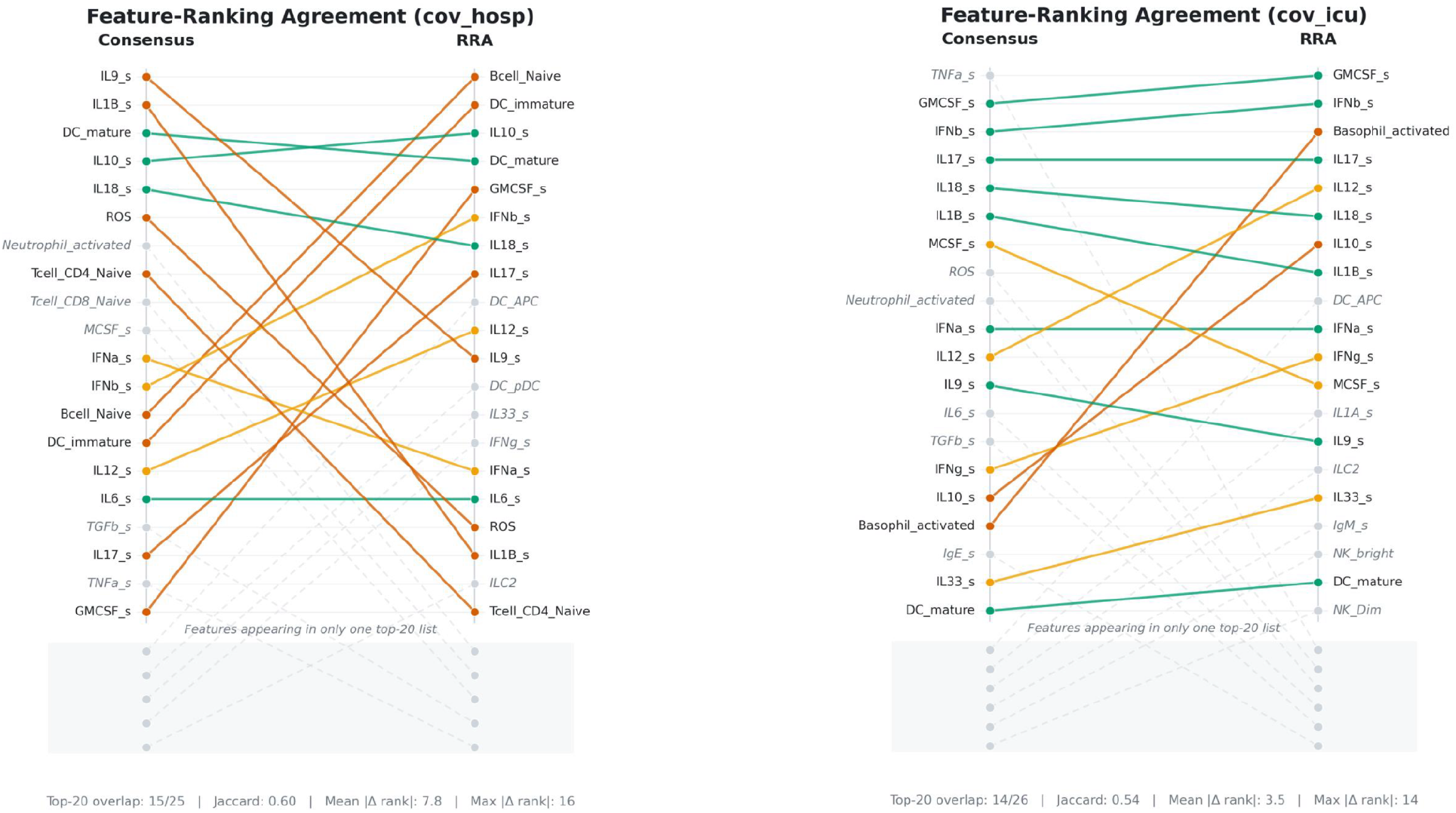
Cross-method agreement of top-ranked features in COV-HOSP and COV-ICU. Top-20 features under the Consensus ranking (left, anchor) and their corresponding ranks under Robust Rank Aggregation (right) for COV-HOSP (**A**,**B**) and COV-ICU (**C**,**D**). Consensus bars encode the per-feature weighted selection score (Σ selection_frequency × mean F1 across all 25 Selector × Classifier pipelines); RRA bars encode −log_10_(ρ_Bonferroni). Solid colors (green, orange, and red): feature in both methods’ top-20 (15/20 for hospitalization and 14/20 in ICU admission).

## METHODS

We present a disease-agnostic machine learning framework that predicts the most significant biomarkers that “define” each of the disease’s two phases (hospitalized vs ICU). The design is intended to be adaptive in the way that the algorithms are not hardwired. Rather, using carefully chosen representatives from each family of classification algorithms to perform classification for an already labeled dataset of patient records as ‘cleared’ and ‘not cleared’. Combined with the most common feature selection algorithms, each classifier predicts the previously assigned label. Based on the performance of the feature selection and classification tasks, we further analyze the top-k performing algorithms for the most dominant features across all the runs.

Depending on the phase (hospitalized vs ICU), and the dataset, the feature selection and classification algorithms will perform differently. For example, SelectKBest+NaiveBayes may be top performing algorithms for COV-HOSP while they may not offer the same value for COV-ICU. Therefore, each prediction must be processed with a mindset of an undefined set of top-k runs until all the experiments are performed and the results are concluded. The most features are then derived from the top runs using a consensus-based heuristic that determines which features are contributing to the “cleared” and which are contributing to the “not cleared” patient condition.

### Feature-selection methods

We selected five feature-selection algorithms to explore the feature space independently, each chosen to capture a different statistical or geometric structure in the data. All methods were configured to retain the top measure features list as follows: (1) ***SelectKBest:*** a univariate ranking that retains the top scoring features, (2) ***Mutual Information:*** Univariate ranking by mutual information between each feature and the target, capturing nonlinear association, (3) Recursive Feature Elimination with a Linear SVM: Iterative elimination of the least-informative feature under a class-balanced linear support vector machine, applied to standardized training data, (4) L1-Regularized Logistic Regression: a sparse logistic regression with class-balanced weights; features with the largest absolute coefficients were retained, (5) ***Variance Inflation Factor (VIF) Filtering:*** Iterative removal of the feature with the highest variance inflation factor, with singular or near-singular features eliminated first, until at most fifteen features remained and all retained features had a VIF below 10.

### Classifiers Representatives

Five classifiers spanning distinct algorithmic families were introduced. (1) ***Logistic Regression:*** serving as a linear baseline, (2) **k-Nearest Neighbors:** Distance-based voting with five neighbors, (3) **Multilayer Perceptron:** A feedforward neural network with a single hidden layer of one hundred units, (4) **Gaussian Naïve Bayes:** a probabilistic classifier using the bayes rule and operates under a Gaussian feature-likelihood assumption, (5) **Decision Tree:** it provides a map that is based on if-else flowchart that supports hierarchical humans reasoning for decision-making. The full design yields twenty-five selector--classifier configurations per run.

### Datasets: The SARS-CoV-2 HOPS vs ICU

We applied the framework to two cohorts derived from a SARS-CoV-2 immune-response dataset: a hospitalization cohort (COV-HOSP) and an intensive-care cohort (COV-ICU). Each record represents a single simulated patient profile characterized by a panel of biological features, including cell-population activations, cytokine concentrations, and intracellular markers. Every record is annotated with respect to infection clearance and assigned a binary label (cleared or not cleared). A measurability annotation accompanies each cohort and indicates which features are considered clinically measurable; only these features were retained for downstream analysis.

The hospitalization cohort comprises 100,000 records and the intensive-care cohort comprises 5,673 records, each described by 108 raw features. After preprocessing, the modeled feature set is reduced to 68 numeric features for COV-HOSP and 65 for COV-ICU. Cohort sizes are summarized in Table 3.

**Table 3.** Tiered discriminatory features per cohort (Consensus rank, RRA rank, Cohen’s *d*, direction)

| Cohort | Feature | Tier | Consensus Rank | RRA Rank | Cohen's d | Direction |
| --- | --- | --- | --- | --- | --- | --- |
| COV-HOSP | DC_mature | 1 | 3 | 4 | -1.13 | ↓ negative |
| COV-HOSP | IL10_s | 1 | 4 | 3 | 2.04 | ↑ positive |
| COV-HOSP | IL18_s | 1 | 5 | 7 | -1.16 | ↓ negative |
| COV-HOSP | IL9_s | 2 | 1 | 11 | 2.90 | ↑ positive |
| COV-HOSP | IFNa_s | 2 | 11 | 15 | 1.50 | ↑ positive |
| COV-HOSP | IFNb_s | 2 | 12 | 6 | 1.11 | ↑ positive |
| COV-HOSP | Bcell_Naive | 2 | 13 | 1 | -1.43 | ↓ negative |
| COV-HOSP | DC_immatu<br>re | 2 | 14 | 2 | -1.45 | ↓ negative |
| COV-HOSP | IL12_s | 2 | 15 | 10 | 1.42 | ↑ positive |
| COV-HOSP | IL6_s | 2 | 16 | 16 | 1.31 | ↑ positive |
| COV-HOSP | IL17_s | 2 | 18 | 8 | -1.16 | ↓ negative |
| COV-HOSP | GMCSF_s | 2 | 20 | 5 | -1.17 | ↓ negative |
| COV-HOSP | DC_APC | 3 | out of top<br>20 | 9 | 0.04 | ↑ positive |
| COV-HOSP | DC_pDC | 3 | out of top<br>20 | 12 | -0.65 | ↓ negative |
| COV-HOSP | IL33_s | 3 | out of top<br>20 | 13 | -1.02 | ↓ negative |
| COV-HOSP | IFNg_s | 3 | out of top<br>20 | 14 | -0.79 | ↓ negative |
| COV-ICU | GMCSF_s | 1 | 2 | 1 | -0.99 | ↓ negative |
| COV-ICU | IFNb_s | 1 | 3 | 2 | 2.57 | ↑ positive |
| COV-ICU | IL17_s | 1 | 4 | 4 | -0.98 | ↓ negative |
| COV-ICU | IL18_s | 1 | 5 | 6 | -0.98 | ↓ negative |
| COV-ICU | IL1B_s | 1 | 6 | 8 | 2.68 | ↑ positive |
| COV-ICU | IFNa_s | 1 | 10 | 10 | 2.63 | ↑ positive |
| COV-ICU | MCSF_s | 2 | 7 | 12 | -0.98 | ↓ negative |
| COV-ICU | IL12_s | 2 | 11 | 5 | 1.46 | ↑ positive |
| COV-ICU | IL9_s | 2 | 12 | 14 | 2.37 | ↑ positive |
| COV-ICU | IFNg_s | 2 | 15 | 11 | -0.93 | ↓ negative |
| COV-ICU | IL10_s | 2 | 16 | 7 | 1.14 | ↑ positive |
| COV-ICU | Basophil_ac<br>tivated | 2 | 17 | 3 | -0.69 | ↓ negative |
| COV-ICU | DC_APC | 3 | out of top<br>20 | 9 | -0.58 | ↓ negative |
| COV-ICU | IL1A_s | 3 | out of top<br>20 | 13 | -0.08 | ↓ negative |
| COV-ICU | ILC2 | 3 | out of top<br>20 | 15 | -0.58 | ↓ negative |

**Table 3.** Cohort sizes (raw, before preprocessing).

| Cohort | Disease | Phase | Records | Raw features | Modeled features after preprocessing |
| --- | --- | --- | --- | --- | --- |
| COV-HOSP | SARS-CoV-2 | Hospitalization | 100,000 | 108 | 68 |
| COV-ICU | SARS-CoV-2 | Intensive care | 5,673 | 108 | 65 |

### PREPROCESSING

For each cohort, the column set was restricted to the intersection of the measurability whitelist and the target column. A curated set of auxiliary columns encoding internal model state, including any feature designated as resting-state, was removed. Target labels were normalized and mapped to a binary encoding, with 0 denoting cleared and 1 denoting not cleared; records carrying unmapped target values were excluded explicitly rather than dropped silently. Constant columns, non-numeric columns, and columns with entirely missing values were then removed, and any remaining record containing a missing predictor value was excluded. No imputation was performed.

The framework assumes independent and identically distributed observations, which is consistent with the simulated provenance of the data. Cohorts exhibiting patient-level grouping or temporal structure would require an alternative cross-validation strategy. **Algorithm 1** Describe the details necessary to process the input data to push it forward into the feature selection and classification stage.

#### Algorithm 1

For Dataset Preprocessing for Cross-Validated Feature Selection

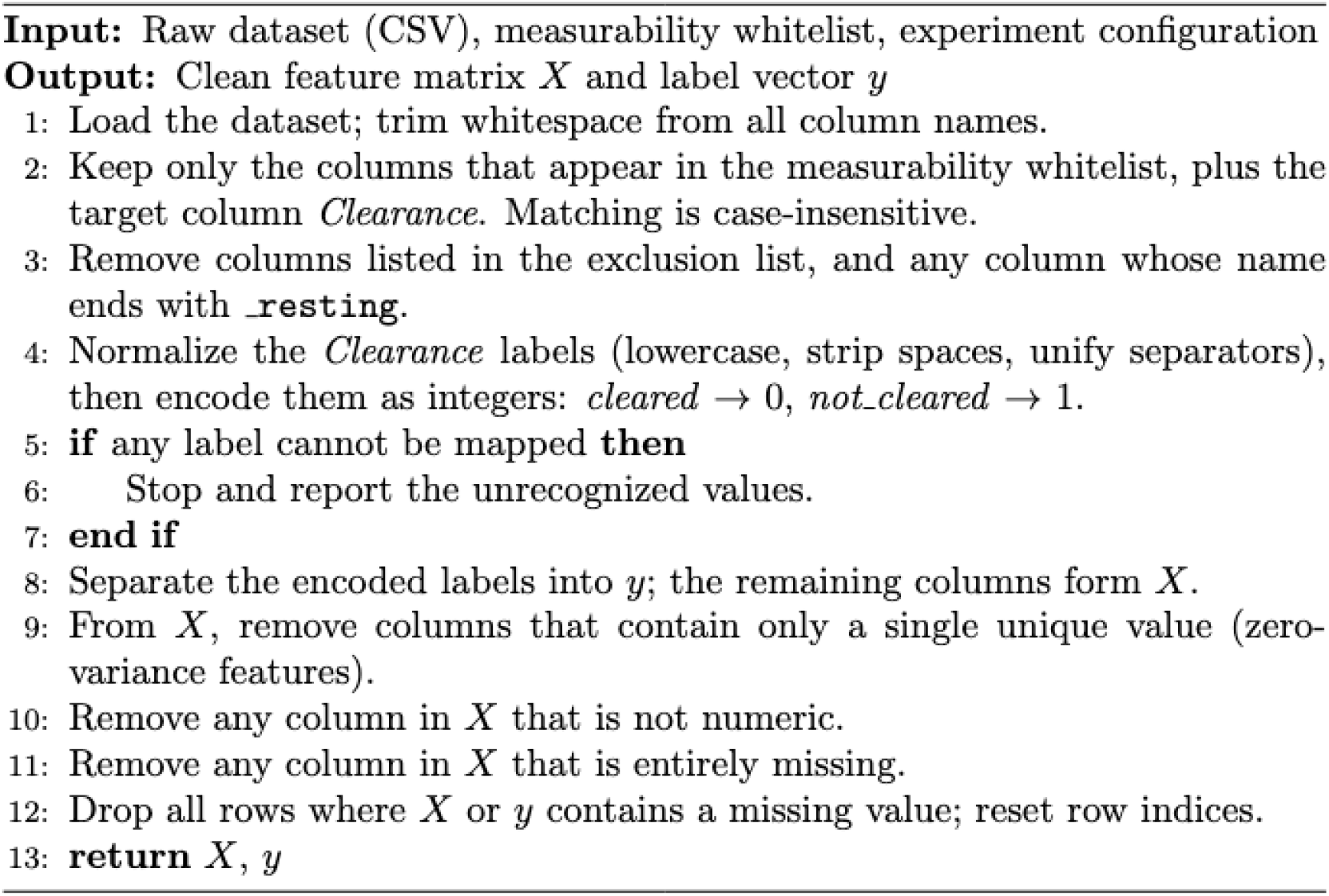

### Cross-validation design

Each combination of feature-selection method and classifier was evaluated under stratified five-fold cross-validation with a fixed random seed to support reproducibility. Feature selection was fitted exclusively on the training fold and applied to the corresponding held-out validation fold, preventing selection leakage. Predictors were standardized within each fold, and classifiers were reinitialized for each split so that no information from earlier folds was carried forward.

### Per-fold metrics and per-run aggregation

For each fold we computed the two-by-two confusion matrix and derived the accuracy, macro-averaged precision, macro-averaged recall, and macro-averaged F1 score. Macro averaging gives equal weight to the cleared and not-cleared classes, which is important given the substantial class imbalance in the hospitalization cohort (approximately 16.6:1) while the ICU imbalance (∼4.5 : 1). For each configuration we report the mean and standard deviation of each metric across the five folds, together with the totals of true and false positives and negatives pooled across folds. **Algorithm 2** shows the feature selection, training, and classification part of the algorithm until a label is produced.

#### Algorithm 2

Shows the repeated process of feature selection combined with classification for each pair of the algorithms to produce the F1 scores and rank the results

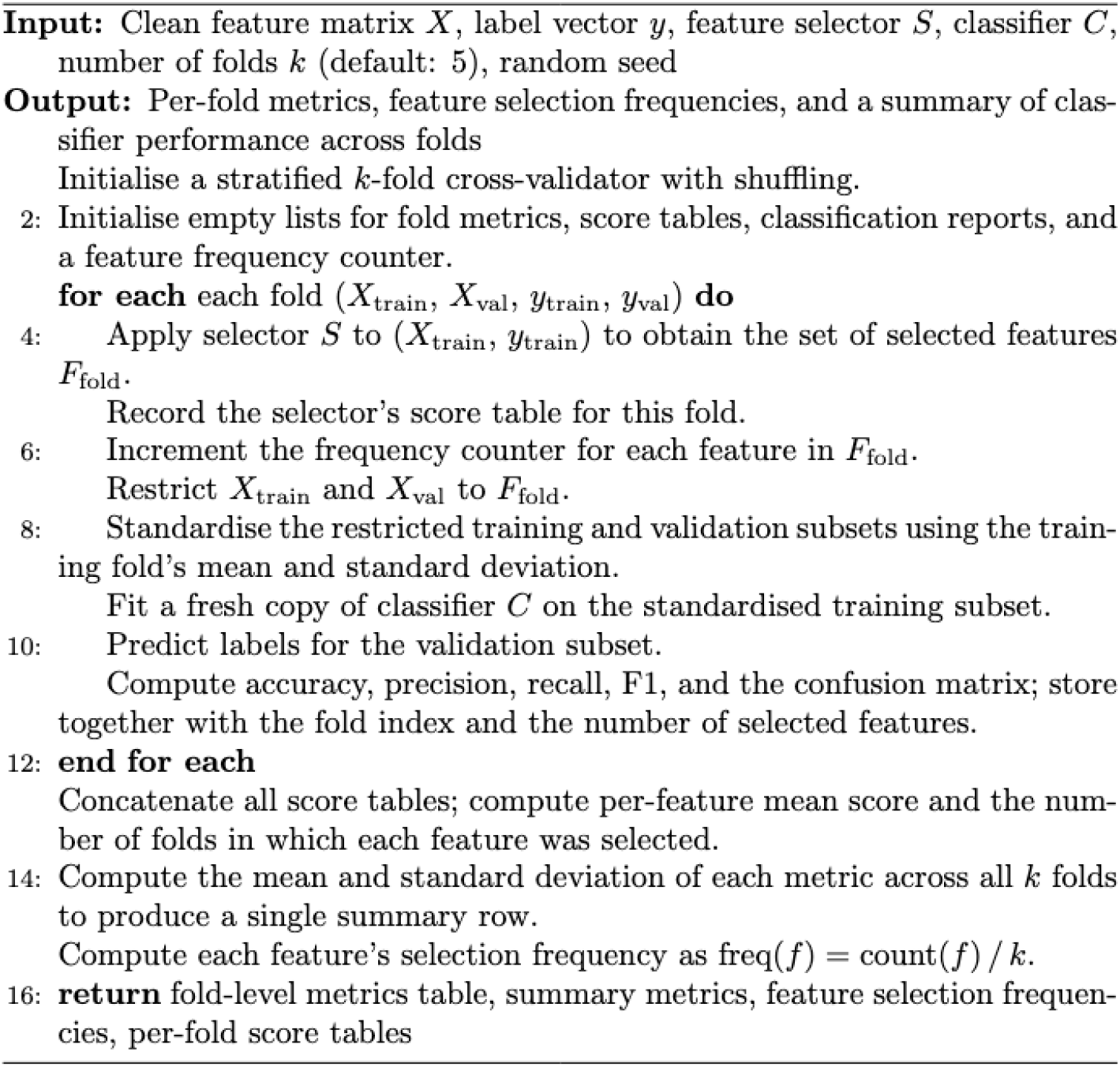

### Cross-run aggregation

Three aggregations were performed across the twenty-five configurations of a single experiment.

1. **Ranked model summary**. All configurations were ordered by mean F1, with ties broken successively by F1 standard deviation, macro recall, macro precision, and accuracy.
2. **Consensus feature ranking**. For each feature, a weighted selection score was computed across all feature selector-classifier combinations as the sum of the product of its within-run selection frequency and the run’s mean F1 score. This ensures that features consistently chosen by higher-performing models contribute more to the final ranking. The full procedure is formalized in **Algorithm 3**, which shows the computational steps necessary to reproduce the ranking score based on the consensus heuristic.
3. **Robust Rank Aggregation (RRA)**. While the weighted selection score ranks features by aggregate importance, it does not account for whether consistent high ranking could arise by chance. Following Kolde et al.^20^, RRA addresses this by treating each selector–classifier run as an independent ranked list. Each run produced a per-feature ranking with normalized ranks in the unit interval. For each feature, we computed the beta-distribution-based order-statistic tail probability across its sorted normalized ranks, took the minimum across rank positions as the raw RRA score, and applied Bonferroni correction across contributing runs. Features were ordered by this score (lower is more significant); for reporting, we used the negative base-ten logarithm of the corrected score, with values above 1.3 indicating p < 0.05 and above 2.0 indicating p < 0.01. Together with the weighted selection score, this approach identifies features that are both performance-weighted and statistically robust consensus biomarkers. **Algorithm 4**. describes the computational steps for producing the RRA score, which is used alongside the weighted selection score to identify the features that most consistently and robustly distinguish the hospitalized from the ICU cohort.

#### Algorithm 3

show the computation steps used to produce a ranking score based on a consensus heuristic driven from the experiment results

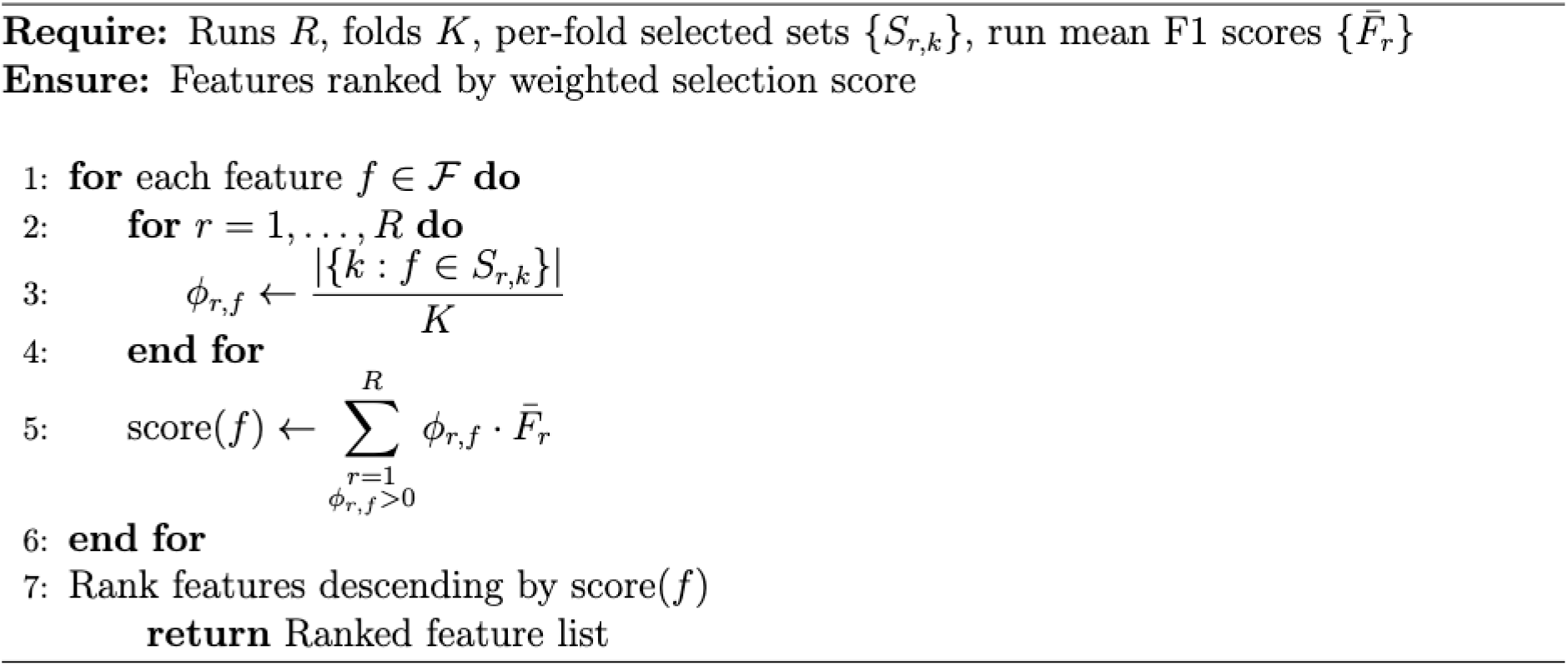

#### Algorithm 4

Robust Rank Aggregation (RRA) aggregates the per-run feature rankings into a single consensus list by testing whether each feature’s rank positions across selector-classifier runs are significantly better than expected by chance, following the order-statistic-based formulation of Kolde et al.

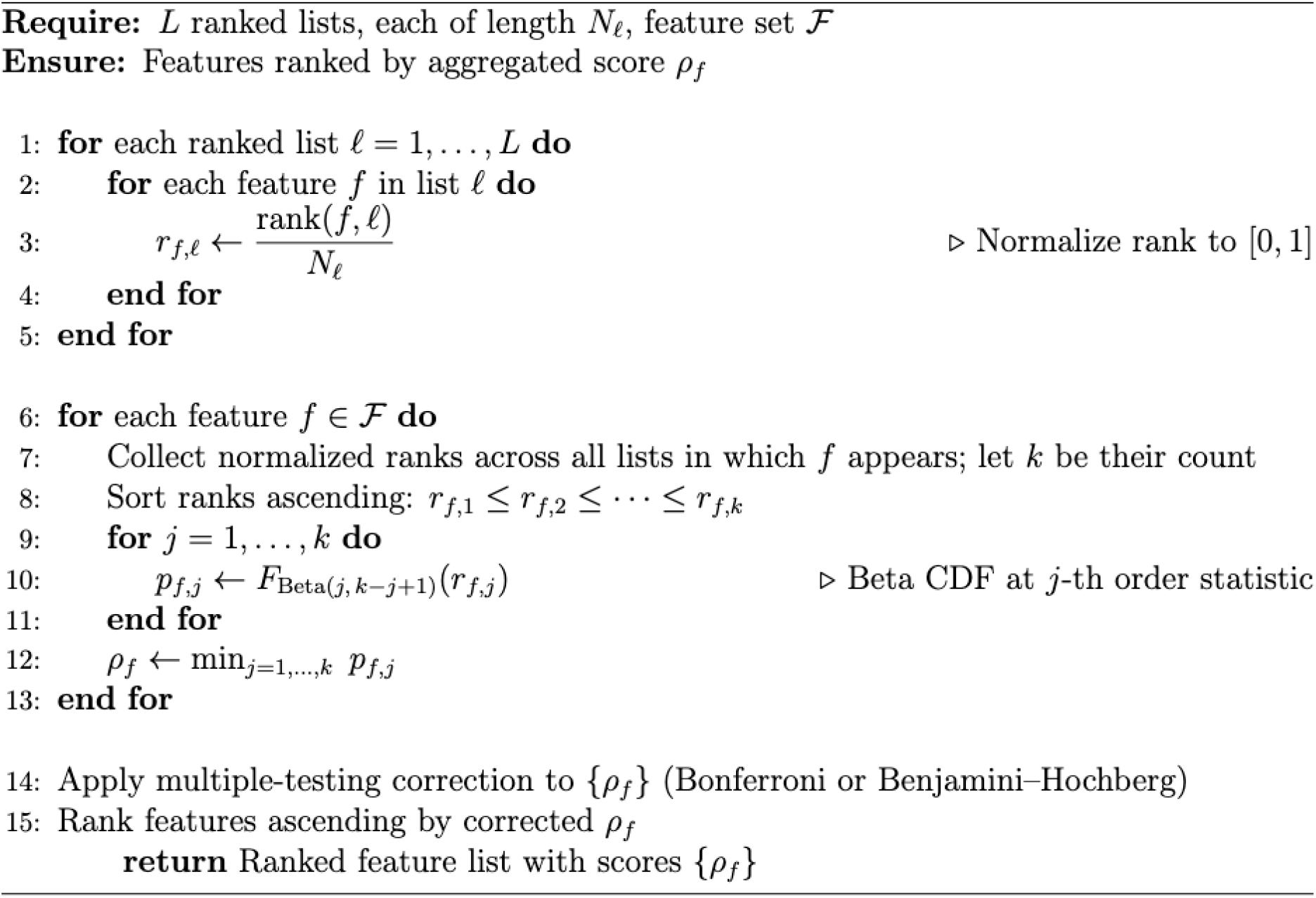

### Discriminatory-biomarker tiers

To translate the rankings into a clinically interpretable summary, we partitioned the union of the top-twenty features under the Consensus and RRA rankings into three hierarchical tiers.

- **Level 1: Strong confidence biomarker**. The feature appears in the top twenty of both rankings, has an absolute rank shift of at most two positions between rankings, and an RRA score of at least 2.0.
- **Level 2: Moderate confidence biomarker**. The feature appears in the top twenty of both rankings but fails one of the Level 1 criteria, either by having a larger rank shift or a weaker RRA score.
- **Level 3: Exploratory confidence biomarker**. The feature appears in the top twenty of only one of the two rankings.

Tier thresholds were chosen a priori and held fixed across cohorts to support sensitivity analyses and cross-cohort comparison. **Algorithm 5** formalizes the tier assignment rules as a function of rank overlap, rank shift, and RRA score.

#### Algorithm 5

Hierarchical Tier Assignment of Consensus Discriminatory Biomarkers

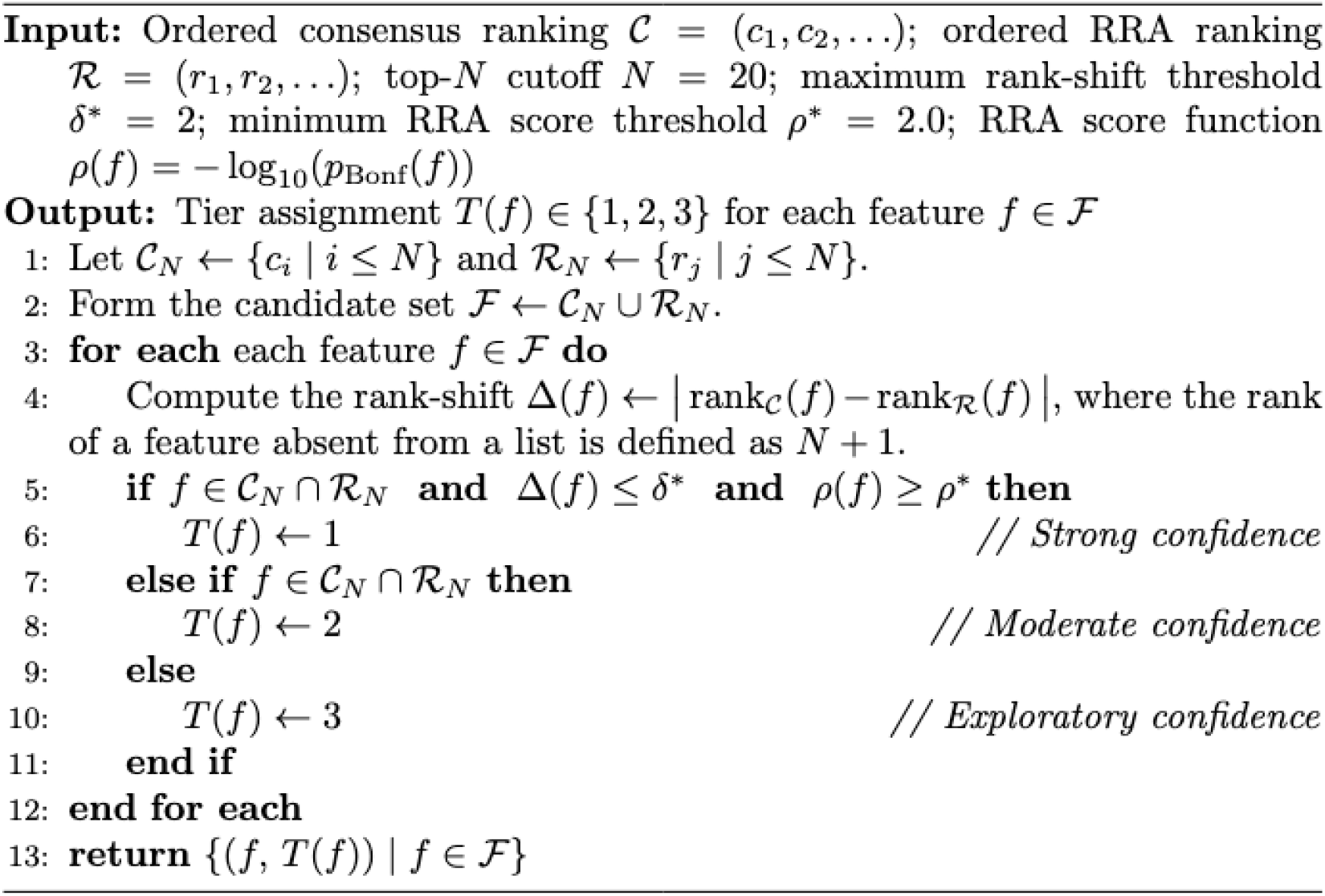

## DISCUSSION

Dysregulated immune responses are hallmarks of disease and disease progression. Here, we describe predicted immune biomarkers signaling an increased risk of hospitalization for COVID-19 patients. Acute viral infection is classically associated with early antiviral signaling followed by inflammation and antigen presentation. In general, airway disease, including asthma, COPD, viral pneumonia, as well as cytokine-driven lung injury in general and COVID specifically, are characterized by changes in cytokines and immune cell numbers and activation states. Elevation of IL-10 (the first tier 1 indicator of hospitalization risk here, Figure 5A) is a classic inflammatory marker across airway disease and has been shown in multiple cohorts to be predictive of COVID disease progression^21–28^. Indeed, IL-10 and IL-6 upregulation (both predicted here) represent some of the most consistently replicated biomarkers of COVID hospitalization across cohorts^21–25,28–32^. Both IL-6 and IL-10 have also been associated with worse clinical outcomes, including transfer to ICU and mortality^21–26,28,29,31–35^. We also identify IL-10 as a tier 2 predictor of ICU transfer (Figure 5B). Similarly, dysfunction in dendritic cells, including a decrease in immature and mature DCs (tier 2 and tier 1, respectively, Figure 5A and a decrease in antigen presenting DCs, tier 3, Figure 5B) are common across severe viral infection. Together, these are suggestive of a general severe airway inflammation signature. More specific to COVID are the hallmarks of type I interferon dysregulation (including increased IFNa and IFNb, tier 2 (Figure 5B) and IFNb, tier 1 (Figure 5A) which are indicative of chronic viral infection^24,31,36^. While severe COVID-19 is characterized first by the “cytokine storm” profile, then by delayed, blunted, or compartmentalized type I IFN activity^25,30,35–38^.

**Figure 5.**
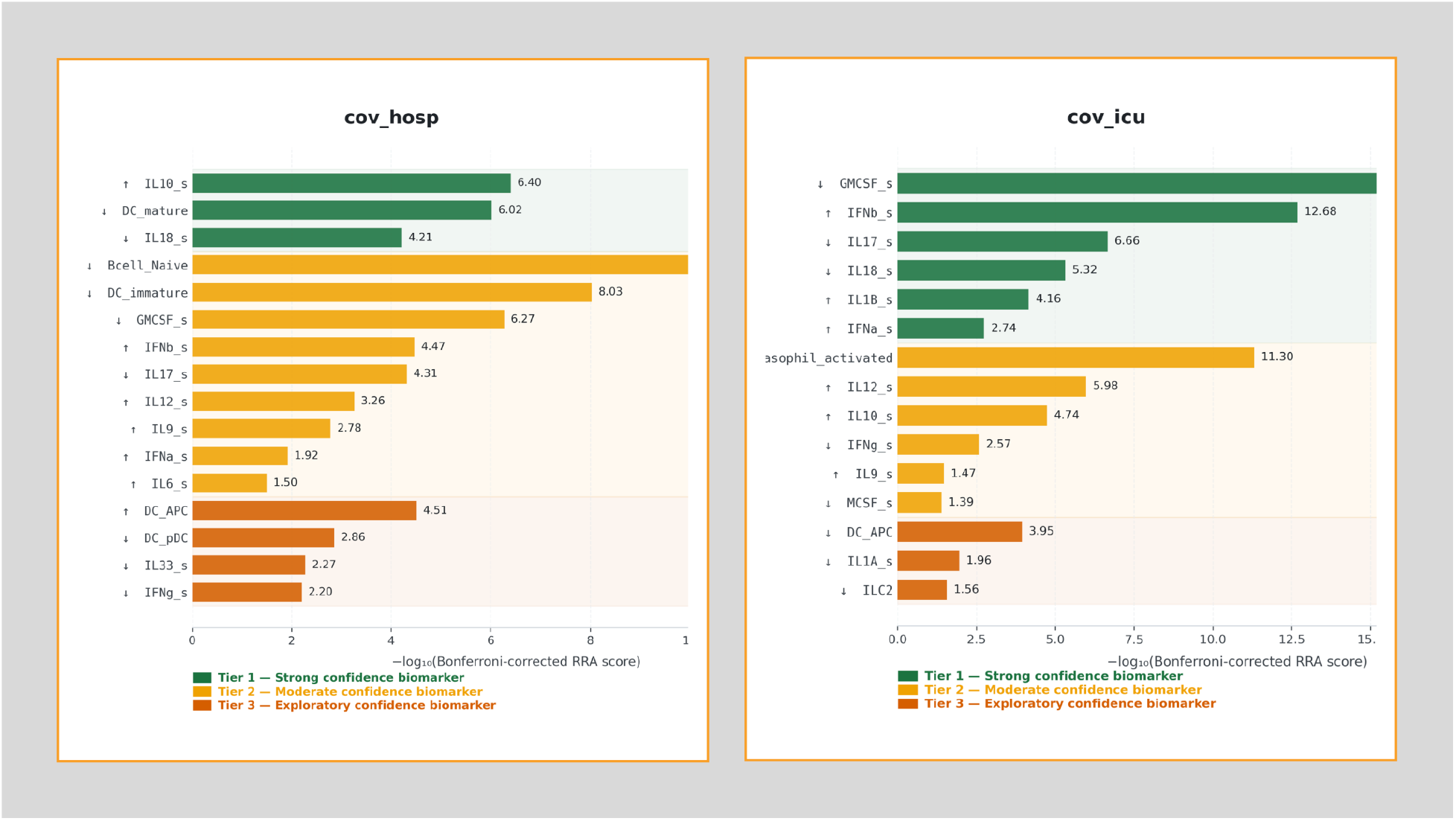
Cohen’s d effect sizes for tiered biomarkers in COV-HOSP and COV-ICU. Tier-wise Cohen’s d values show strong, concordant effect directions across cohorts. The largest Tier-1 effects were IL10_s, IL18_s, DC_mature in COV-HOSP and IL1B_s, IFNa_s, IFNb_s in COV-ICU. Ten markers appeared in both cohorts, with 9/10 sharing the same direction; the lone discrepancy (DC_APC) was negligible in magnitude. All five cross-phase candidates (IL18_s, IL17_s, IFNa_s, IFNb_s, GMCSF_s) showed consistent signs across cohorts, supporting their robustness. Table 2 lists the top features, consensus/RRA scores, and effect directions.

Additionally, dysfunction of antigen presenting cells predicted here through decreased DC subsets as well as decreased naïve B-cells (tier 2), which are necessary for long-term immunity via their expansion to mature and plasma B cells^38^. While antigen presenting cells may rise transiently with infection, severe disease is associated with the loss of mature DCs^38^ and impaired maturation or costimulatory capacity, consistent with defective T-cell priming and weakened adaptive immunity. Lymphopenia is a predictor of disease severity both in COVID (including for ARDS development, the need for ICU transfer, and mortality)^24,25,27,29,32–37,39^. Several of the cytokines predicted in tier 2 as increased in individuals at risk for hospitalization (Figure X) also support this, as both IL-12 (generally considered a Th1-promoting cytokine which increases in severe inflammation) and IL-9 (implicated in severe disease). IL-1B, INFa/b, IL-10, IL-9, and IL-12 have also been linked to critical illness or ICU admission in at least some cohorts of patients (and are often not all tested in parallel or uniformly across cohorts, perhaps contributing to the variation seen in clinical samples)^22,25,26,33–35^. IL-1B and IL-10, specifically, are already currently used in prognostic ratios and ICU-specific biomarker panels, supporting the validity of this model^22,25,34^. Biologically, the specificity of this predicted response points more to a dysregulation of immune control rather than global cytokine excess. This is supported by the downregulation of GM-CSF (top tier 1 predictor of ICU (Figure 5B) and tier 2 (Figure 5A)) and M-CSF (tier 2 Figure 5B), both of which are central to monocyte/macrophage and myeloid cell development and function, and where reduced levels would indicate exhausted or dysregulated myelopoiesis. Biologically, this suggests impaired antigen presentation and weaker T-cell priming, both expected and seen in sicker patients. Reduced IL-17, IL-18, and IFNg (Figure 5B) may also indicate that on the path to ICU-level disease, the immune response is no longer productively antiviral but instead becomes fragmented and functionally suppressed in key compartments. This is also supported by lower IL-1a and ILC2, which may be readouts for tissue-immune disruption^36^; e.g., combined as signatures of hyperinflammation with a collapse of normal immune regulation and antigen presentation.

## FUTURE DIRECTION

Moving forward, we intend to employ the MarkerScout pipeline to identify critical immune-response biomarkers associated with additional pathologies, specifically influenza and malaria . This future research will involve analyzing outcomes from cohorts of patients who required hospitalization or intensive care unit admission.

## ACKNOWLEDGMENTS

This work was supported by the National Institutes of Health under Grant No. #R35GM119770, and the University of Nebraska-Lincoln Grand Challenges Catalyst Award to Tomáš Helikar.

We are thankful to Skylar Loecker for their valuable discussions during the methodology and investigation phases of this project.

## AUTHOR CONTRIBUTIONS

Conceptualization, T.H; methodology T.H, F.A.N, and A.A.H.; Investigation, F.A.N, R.M, P.P, L.B.C, A.A.H, and T.H; writing – original draft, A.A.H; writing review & editing, A.A.H, L.B.C, R.M, and T.H; funding acquisition, T.H; supervision, A.A.H, T.H.

## DECLARATION OF INTERESTS

T.H is a founder and shareholder of Discovery Collective, Inc., and ImmuNovus, Inc.

## DECLARATION OF GENERATIVE AI AND AI-ASSISTED TECHNOLOGIES IN THE WRITING PROCESS

Claude (Anthropic) and builtin Gemini in Google Docs were used to reformat and restructure these outputs into manuscript-ready sections. All AI-generated content was reviewed, edited, and validated by the authors, who take full responsibility for the scientific content.

